# To what extent gene connectivity within co-expression network matters for phenotype prediction?

**DOI:** 10.1101/523365

**Authors:** Aurélien Chateigner, Marie-Claude Lesage-Descauses, Odile Rogier, Véronique Jorge, Jean-Charles Leplé, Véronique Brunaud, Christine Paysant-Le Roux, Ludivine Soubigou-Taconnat, Marie-Laure Martin-Magniette, Leopoldo Sanchez, Vincent Segura

## Abstract

Recent literature on the differential role of genes within networks distinguishes core from peripheral genes. If previous works have shown contrasting features between them, whether such categorization matters for phenotype prediction remains to be studied. We sequenced RNA in a *Populus nigra* collection and built co-expression networks to define core and peripheral genes. We found that cores were more differentiated between populations than peripherals while being less variable, suggesting that they have been constrained through potentially divergent selection. We also showed that while cores were overrepresented in a subset of genes deemed important for trait prediction, they did not systematically predict better than peripherals or even random genes. Our work is the first attempt to assess the importance of co-expression network connectivity in phenotype prediction. While highly connected core genes appear to be important, they do not bear enough information to systematically predict better quantitative traits than other gene sets.

## Introduction

Gene-to-gene interaction is a pervasive although elusive phenomenon underlying phenotype expression. Genes operate within networks with more or less mediated actions on the phenome. Systems biology approaches are required to grasp the functional topology of these networks and ultimately gain insights into how gene interactions interplay at different biological levels to produce global phenotypes (Mackay et al., 2009). New sources of information and their subsequent use in the inference of gene networks are populating the wide gap existing between phenotypes and DNA sequences and, therefore, opening the door to systems biology approaches for the development of context-dependent phenotypic predictions. RNA sequencing (RNAseq) is one of such new sources of information that can be used to infer gene networks (Han et al., 2015).

Among the many works on gene network inference based on transcriptomic data, two recent studies aimed at characterizing the different gene roles within co-expression networks (Josephs et al., 2017; Mähler et al., 2017). Josephs et al. (2017) studied the link between gene expression, gene connectivity (Langfelder and Horvath, 2008), divergence (Williamson et al., 2005) and traces of natural selection (Josephs et al., 2015; Sicard et al., 2015) in a natural population of the plant *Capsella grandiflora*. They showed that both connectivity and local regulatory variation on the genome are important factors, while not being able to disentangle which of them is directly responsible for patterns of selection among genes. Mähler et al. (2017) recalled the importance of studying the general features of biological networks in natural populations. With a genome-wide association study (GWAS) on expression data from RNAseq, they suggested that purifying selection is the main mechanism maintaining functional connectivity of core genes in a network and that this connectivity is inversely related to eQTLs effect size. These two studies start to outline the first elements of a gene network theory based on connectivity, stating that core genes, which are highly connected, are each of high importance, and thus highly constrained by selection. In contrast to these central genes, there are peripheral, less connected genes, never far from a core hub. These peripheral genes are less constrained than core genes and consequently, they harbor larger amounts of variation at population levels.

Furthermore, classic studies of molecular evolution in biological pathways can help us understand the link between gene connectivity and traits. Several articles showed that selection pressure is correlated to the gene position within the pathway, either positively (Han et al., 2013; Lu, 2003; Rausher et al., 2008, 1999; Riley et al., 2003; Yu et al., 2011) or negatively (Han et al., 2013; Jovelin and Phillips, 2011; Song et al., 2012; Wu et al., 2010), depending on the pathway. Jovelin and Phillips (2011) showed that selective constraints are positively correlated to expression level, confirming previous studies (Drummond et al., 2005; Duret and Mouchiroud, 2000; Pál et al., 2001). Montanucci et al. (2011) showed a positive correlation between selective constraints and connectivity, although such a possibility remained contentious in previous works (Bloom and Adami, 2004; Fraser and Hirsh, 2004).

While Josephs’ (Josephs et al., 2017) and Mahler’s (Mähler et al., 2017) studies framed a general view of genes organization based on topological features described in molecular evolution studies of biological pathways, a point remains quite unclear so far: to what extent core and peripheral genes based on connectivity within a co-expression network are involved in the definition of a phenotype? One way to clarify this would be to study the respective roles of core and peripheral genes, as defined on the basis of their connectivity within a co-expression network, in the prediction of a phenotype. Even if predictions are still one step before validation by in vivo experiments, they already represent a landmark that may not only be correlative but also closer to causation, depending on the modeling strategy.

Present study aims at exploring gene ability to predict traits, with datasets representing core genes and peripheral genes. By making use of two methods to predict these phenotypes, a classic additive linear model, and a more complex and interactive neural network model, we further aimed at studying the mode of action of each type of genes, in order to gain insight into the genetic architecture of complex traits. On the one hand, genes that are better predictors with an additive model are supposed to have an overall more additive, direct mode of action representing a gene that would be downstream in a biological pathway. We expect core genes to display such additive behavior, with a high but selectively constrained expression level (Jovelin and Phillips, 2011; Montanucci et al., 2011). On the other hand, genes being better predictors with an interactive model are supposed to be upstream in pathways. We expect peripheral genes to behave interactively, with a lower but relatively more variable expression level. With a lower variation, we also expect core genes to be worse predictors for traits than peripheral genes unless the former also bear larger effects.

To answer the questions concerning the respective roles of core and peripheral genes on phenotypic variation, we have sequenced the RNA of 459 samples of black poplar (*Populus nigra*), corresponding to 241 genotypes, from 11 populations representing the natural distribution of the species across Western Europe. We also have for each of these trees phenotypic records for 17 traits, covering the growth, phenology, physical and chemical properties of wood. They cover two different environments where the trees were grown in common gardens, in central France and northern Italy. With the transcriptomic data, we built a co-expression network in order to define contrasting gene sets according to their connectivity within the network. We then asked whether these contrasting sets differed in terms of both population and quantitative genetics parameters and quantitative trait prediction.

## Results

### Wood samples, phenotypes, and transcriptomes

Wood collection and phenotypic data (**Table S1**) have been previously described (Gebreselassie et al., 2017). Further details are provided in the materials and methods section. The complete pipeline is sketched in **Figure 1**. Briefly, we are focusing on 241 genotypes coming from different natural populations in western Europe and planted in 2 common gardens (to avoid the confounding between genetic and large environmental effects) at two different locations in 6 replicated and randomized complete blocks, in Orléans (central France) and Savigliano (northern Italy). A total of 17 phenotypic traits (**Table S1**) have been collected on these genotypes (10 traits in Orléans and 7 in Savigliano). In Orléans only, we used 2 clonal trees per genotype (from 2 blocks) to sample xylem and cambium during the 2015 growing season for RNA sequencing. No tree from Savigliano was used for RNAseq. Because of sampling and experimental mistakes that were further revealed by the polymorphisms in the RNA sequences, we ended up with 459 samples for which we confirmed the genotype identity (comparison to previously available genotyping data from an SNP chip). These samples correspond to 218 genotypes with two biological replicates and 23 genotypes with a single biological replicate. We mapped the sequencing reads on the *Populus trichocarpa* transcriptome (v3.0) to obtain gene expression data.

**Figure 1:**
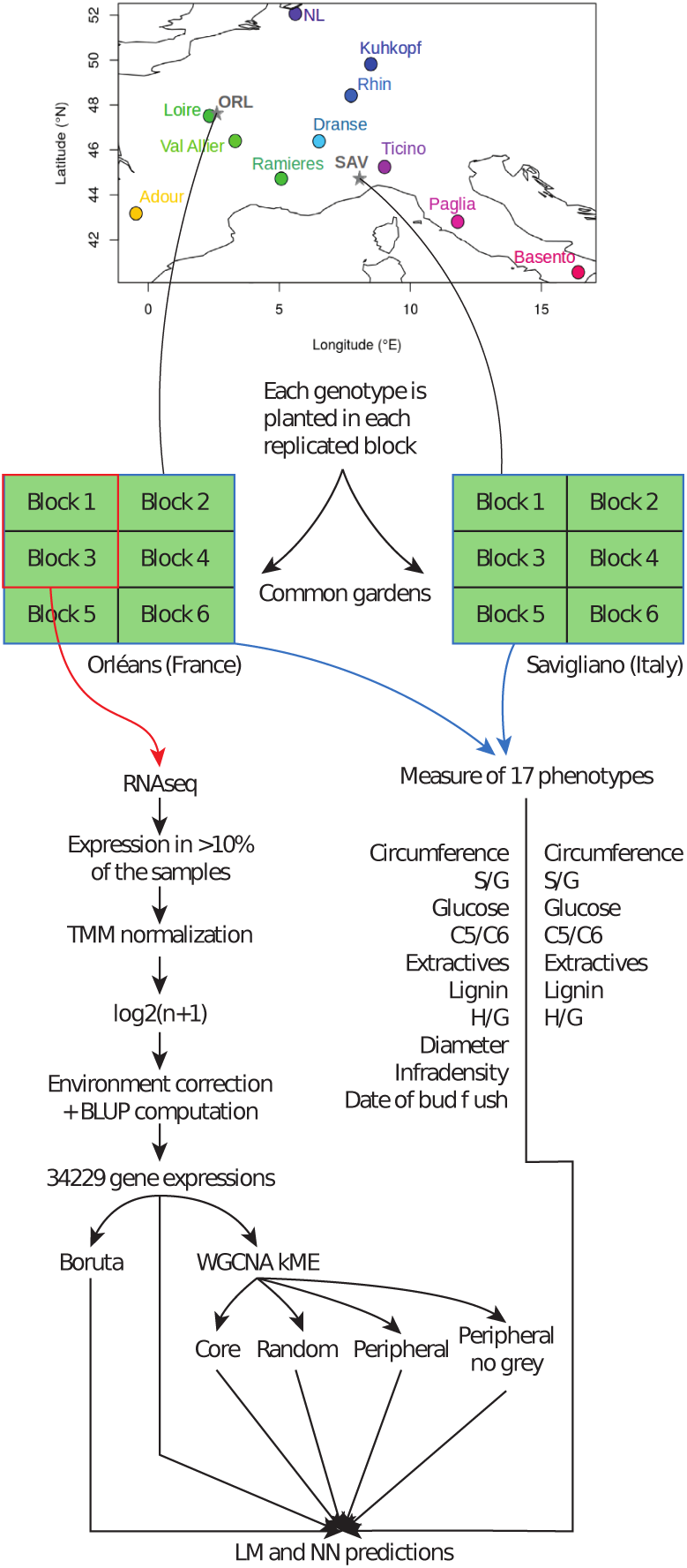
General sketch of the experiment. From the top to the bottom: Map of the location of the different populations sampled for this experiment. From these populations, genotypes were collected and planted in 2 locations (Orléans, in central France, and Savigliano, in northern Italy). At each site, we planted 6 clones of each genotype, 1 in each of the 6 blocks, and their position in each block was randomized. For all the blocks, we collected phenotypes: 10 in Orléans (circumference, S/G, glucose, C5/C6, extractives, lignin, H/G, diameter, infradensity and date of bud flush) and 7 in Savigliano (circumference, S/G, glucose, C5/C6, extractives, lignin, H/G). Only on the clones of 2 blocks in Orléans, we performed the RNA sequencing and treatment of data. The treated RNAseq data were used with different algorithms and in different sets to predict the phenotypes measured on the same trees (in Orléans) or on the same genotype but on different trees (in Savigliano).

We did PCA analyses on the cofactors that were presumably involved in the experience, to look whether any confounding effect could be identified (**Figure S1**). No clear segregation was found for any of those, except for the ones associated with block, date and hour of sampling. We used a linear mixed-model framework to correct the effects of these cofactors on each transcript (see the materials and methods section for a formal description of the model used), with the breedR R package (Muñoz and Sanchez, 2017), and further computed from the models the complete BLUP for each genotype. Hereafter, we refer to this set of BLUPs for the 34,229 transcripts as the full gene set (83% of annotated transcripts).

### Clustering and network construction

The classical approach to build a signed scale-free gene expression network is to use the weighted correlation network analysis (implemented in the WGCNA R package (Langfelder and Horvath, 2008)), using a power function on correlations between gene expressions. We chose to use Spearman’s rank correlation to avoid any assumption on the linearity of relationships. The scale-free topology fitting index (*R*^2^) did not reach the soft-threshold of 0.85, so we chose the classical power value of 12, corresponding to the first decrease in the slope growth of the index, resulting in an average connectivity of 195.2 (**Figure 2A**). We detected 16 gene expression modules (**Table S2**) with automatic detection (merging threshold: 0.25, minimum module size: 30, **Figure 2B**). Spearman correlations between phenotypic and expression data, presented in the lower panel of **Figure 2B** below the module membership of each gene, display a structure when the order follows the gene expression tree. The traits themselves are line ordered according to clustering on their scaled values to represent their relationships (**Figure S2**). Interestingly, some patterns in the correlation between expression and traits do not follow what we would expect from the similarity between traits (5 traits out of 7 with data in both geographical sites). For instance, in the group composed of S/G ratios and glucose composition, the patterns were more similar between sites across traits than between traits across sites (**Figure 2B, Figure S3**). Complex shared regulations mediated by the environment seem to be in control of these phenotypes, suggesting site-specific genetic control. Otherwise, glucose composition in Savigliano, wood basic density, and extractives in Orléans presented similar patterns, contrarily to what would be expected from the correlations between these traits. These results from the comparative analysis of correlations pinpoint some underlying links between traits that are not obvious from factual phenotypic and genetic correlations between traits.

**Figure 2:**
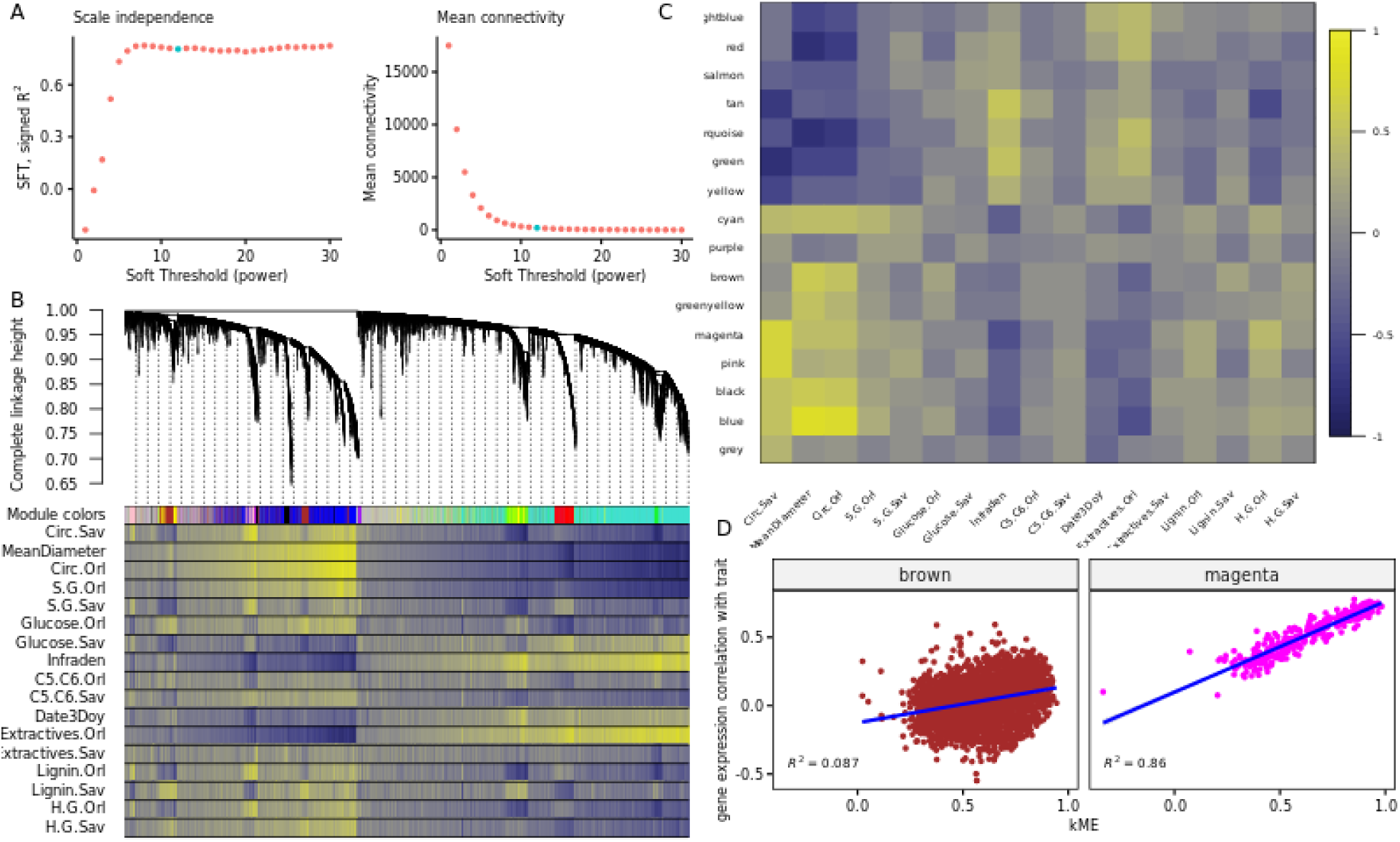
GCNA analysis of gene expression data. (A) Selection of the soft threshold (green dot) based on the correlation maximization with scale-free topology (left panel) producing low mean connectivity (right panel). (B) Gene expression hierarchical clustering dendrogram, based on the Spearman correlations (top panel), resulting in clusters identified by colors (first line of the bottom panel). Spearman correlations between gene expressions and traits values are represented as color bands on the other lines of the bottom panel, from highly negative correlations (dark blue) to highly positive correlations (light yellow), according to the scale displayed in panel C. (C) Spearman correlation between eigengenes (the best theoretical representative of a gene expression module) of modules identified in the previous panel and traits, again on a dark blue (highly negative) to light yellow (highly positive) scale. (D) Focus on two modules from the previous graph, representing the correlations between gene expression correlation with the circumference in Savigliano and centrality in the module. These two panels represent the strongest (right panel, magenta module, *R*^2^ = 0.86) and the weakest (left panel, brown module, *R*^2^ = 0.09) correlations with the corresponding trait.

To get further insight into the relationships between module composition and traits, we looked at the strongest correlations between the best theoretical representative of a gene expression module (eigengene) and each trait, in order to identify genes in relevant modules with an influence on the trait (**Figure 2C**). Following a Bonferroni correction of the p-values provided by WGCNA, only 80 correlations remained significant (*p* ≤ 0.05) out of the initial 272 traits by module combinations. Six traits displayed no significant correlations with any module (Glucose.Sav, both C5.C6, Extractives.Sav, Lignin.Sav and H.G.Sav) and 1 module was not significantly correlated with any of the traits studied (purple, **Figure S3**). In significantly correlated modules, gene expression correlation with trait was also significantly correlated with centrality in the module (represented by the kME, the correlation with the module eigengene), while no correlation was found in poorly correlated modules (**Figure 2D, Figure S4**). In other words, there is a three-way correlation. The genes with the highest kME in a given module are the most correlated to the eigengene and, consequently, are also the most correlated to those traits with the largest correlation with the module eigengene. Although this is somehow expected, it underlines the usefulness of kME as a centrality score to further characterize the genes within each module. We thus used this centrality score to define further the topological position of our gene expressions in the network and to serve as a basis for role comparisons between genes. For each gene, we used its highest absolute score, which corresponds to its score within the module to which it was assigned. We selected the 10% of genes with the highest global absolute scores to define the core genes group, and 10% with the lowest global absolute scores to define the peripheral genes group. Finally, we selected 100 samples of 3422 (10%) random genes as control groups (**Figure S5**, bottom panel).

One particular module from the WGCNA clustering is the grey module. This module typically gathers genes with poor membership to any other module. In our case, it is the 2nd largest module, with 7674 genes (23% of the full set). It gathers the vast majority of genes with very low kME (**Figure S5**, bottom panel) and 99% of the peripheral genes set (**Table S4**).

While it is typically discarded in classic clustering studies, we chose to maintain it and rather understand its composition and role, by adding to the comparative study two peripheral sets, one with and one without grey module genes (subsequently called “peripheral NG”, NG for “no grey”).

To assess the robustness of WGCNA analysis results, we compared it to a k-means clustering (R package coseq, (Rau and Maugis-Rabusseau, 2017)) of the gene expressions (**Figure S6A**). The distribution of WGCNA and k-means’ clusters showed a correlation of −0.49 (**Figure S6B**). k-means clustering tends to form groups of comparable size (Biernacki et al., 2006), which does not seem biologically credible. Furthermore, the correlations between the k-mean modules eigengenes and traits were lower than with WGCNA’s, with a poor repartition of the different modules on the first 2 principal component analysis space (**Figure S6C**). We thus preferred WGCNA clustering to k-means clustering for this analysis.

### Heritability and population differentiation of modules

To get further insights into the biological role of core and peripheral genes at population levels, we looked at the distribution of various characteristics between gene sets (**Figure 3**): gene expression level, several classical population statistics, including heritability (*h*^2^), coefficient of quantitative genetic differentiation (*Q*_*ST*_), coefficient of genetic variation (*CV*_*g*_), gene diversity (*Ht*), and a contemporaneous equivalent to *F*_*ST*_ for genome scans (*PCadapt score*). Gene expression level, *h*^2^, *Q*_*ST*_, and *CV*_*g*_ were computed from gene expression data, while *Ht* and *PCadapt score* (Luu et al., 2017) were computed from polymorphism data (SNP) and averaged per gene model. For more details see the materials and methods section.

**Figure 3:**
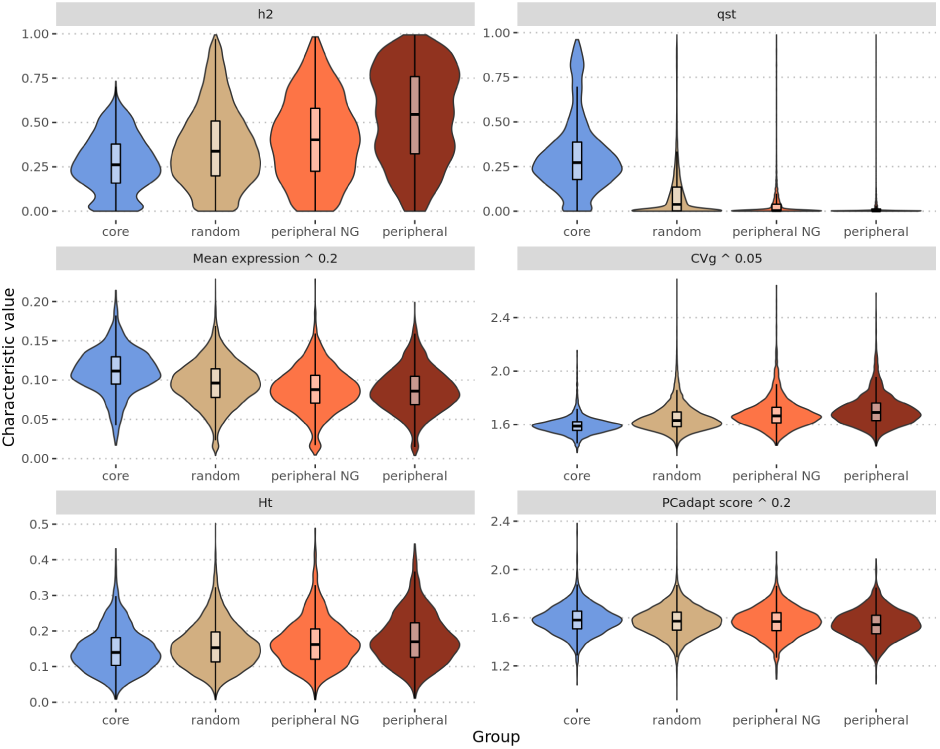
Heritability *h*^2^, differentiation *Q*_*ST*_, gene mean expression (in counts per million, power 0.2), genetic variation coefficient *CV*_*g*_ (power 0.05), overall gene diversity *Ht* and *PCadapt score* (power 0.2) violin and box plots with median (black line) and interquartile range (black box) for each of the core (in blue), random (in grey), peripheral NG (in orange) and peripheral (in brown) gene sets.

Globally, there is a clear trend from core to random, to peripheral NG and to peripheral among these characteristics: with an increase for *h*^2^, *CV*_*g*_ and *Ht*, and a decrease for *Q*_*ST*_, expression and *PCadapt score*. The only differences that are not significant after Bonferroni correction are those between peripheral NG and peripheral sets in gene expression (p-value = 0.14) and between random and peripheral NG sets in the *PCadapt score* (p-value = 0.39). All the other comparisons have p-values below 0.001.

Altogether, these statistics showed clear differences between core and peripheral genes: core genes are highly expressed, highly differentiated between populations in their expression and by their allele frequencies at linked markers, and with generally low levels of genetic variation. Contrastingly, peripheral genes are poorly expressed, poorly differentiated between populations, with generally higher genetic variation.

### Boruta gene expression selection

In addition to previous gene sets building (full, core, random, peripheral NG and peripheral), we wanted to have a set of genes being relevant for their predictability of the phenotype. Our hypothesis here was that this set would be the one that enables the best prediction of a given trait but with a limited gene number. For that purpose, we performed a Boruta (Boruta R package, (Kursa and Rudnicki, 2010)) analysis on 60% of the full genes set (training set). This algorithm performs several random forests to analyze which gene expression profile is important to predict a phenotype. We tested 4 different p-values for this algorithm, as we originally wanted to relax the selection and get eventually sets of different sizes. However, the number of genes selected decreased while relaxing the p-value (613, 593, 578 and 578 respectively for 0.01, 0.05, 0.1 and 0.2). Among the 4 p-values tested, 190 genes were systematically selected (114 are core, 2 are peripheral NG and 2 are peripheral genes), and 153 were selected on 3 of the 4 p-value sets (73 are core, 4 are peripheral NG and 4 are peripheral genes). There is a 6.61 mean over-representation of core genes for the 4 p-values and 0.30 and 0.31 under-representation of respectively peripheral NG and peripheral genes (**Figure S7**). In the end, with a p-value of 0.01, a pool of 613 unique gene expressions was found to be important to predict our phenotypes. Traits with the highest number of important genes are related to growth. For the other traits, we always have more genes selected when the trait is measured in Orléans compared to Savigliano (respective medians of 23 and 10), which fits well with the fact that RNA collection was performed on trees in Orléans. On average, genes that were specific to single traits represented 94% of selected genes, 1 gene was shared across sites for a given trait, genes shared by trait category (growth, phenology, physical, chemical) were 4%, and genes shared among all traits were 2%.

### Phenotype prediction with gene expression

For our 6 genes sets (full, core, random, peripheral NG, peripheral and Boruta), we trained two contrasting classes of models to predict the phenotypes: an additive linear model (ridge regression) and an interactive neural networks model. For the former, we used ridge regression to deal with the fact that for all gene sets the number of predictors was larger than the number of observations. For the latter, we chose neural networks as a contemporary machine-learning method, which is not subjected to dimensionality problems (González-Recio et al., 2014) and is able to capture interactions without a priori explicit declaration between the entries, here gene expressions.

These contrasting models let us capture more efficiently either additivity or interactivity and are thus likely to inform us about the preferential mode of action of each gene set depending on their relative performances in predictability. **Figure 4** and **Figure S8** show that for linear modeling with ridge regression, the best genes set to predict phenotypes was the full set, as expected because it contains more information, followed, more surprisingly, by the peripheral and peripheral NG genes set, then the random, core and Boruta sets (respective mean prediction *R*^2^ across all traits of 0.22, 0.21, 0.20, 0.19, 0.18 and 0.17). On the contrary, for neural network modeling, random genes constituted the worst set by far, followed by core, peripheral, peripheral NG and Boruta sets (respective mean prediction *R*^2^ across all traits of 0.14, 0.16, 0.17, 0.18 and 0.22). We have not been able to compute neural network models with the full set as the number of predictors remains too large to be fitted within a reasonable time on computing clusters. Across phenotypes, predictions were generally slightly less variable under neural networks than under the ridge regression counterpart (interquartile range mean division by 1.12).

**Figure 4:**
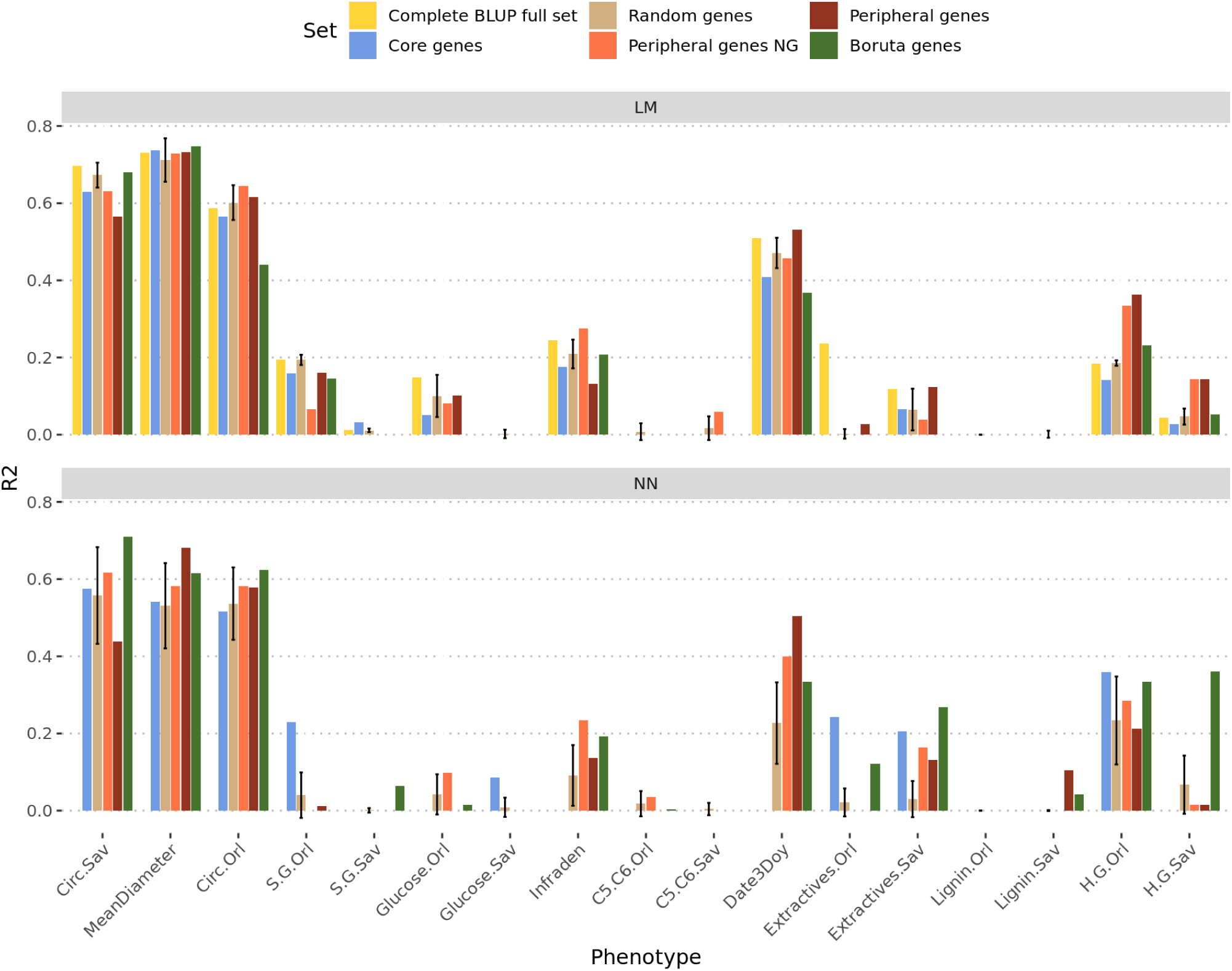
Predictions scores on test sets (*R*^2^ on the y axis) for the 2 algorithms (LM Ridge, top panel; neural network, bottom panel) for each phenotypic trait (on the x axis). The color of each bar represents the gene set that has been used for the prediction. Intervals for the random set represent the first and third quartiles of the distribution of the 100 different realizations, while the height of the bar corresponds to the median.

To further investigate the behavior of genes with different positions in the network with respect to the prediction model used, we computed 2 types of differences: between LM and NN prediction scores for each gene set (**Figure S9A**), and between core and peripheral genes sets for LM and for NN models (**Figure S9B**). As a null reference for inference in the between sets difference (**Figure S9B**), we computed the differences between all the 100 random sets, for a total of 4950 differences corresponding to all pairwise differences, excluding reciprocals and self-comparisons. In the top panel, a positive difference indicates that LM predicted better than NN and *vice versa*, while in the bottom panel, a positive difference indicates an advantage of core genes sets over peripherals and, conversely, a negative difference indicates an advantage of peripheral genes. In any of the two panels, we did not detect any systematic difference, which would have led us to conclude on more interactivity or more additivity for any gene set. Moreover, the few cases where a difference could have been noted are due to very poor prediction scores. The only difference that can be noted is the difference between core and both peripheral genes in NN for the date of bud burst (Date3Doy), in favor of the peripheral genes.

Finally, we investigated to what extent trait *Q*_*ST*_ would influence the prediction scores of each combination of set and algorithm. We thus separated traits according to whether their *Q*_*ST*_ is above or below the 99^*th*^ percentile of the *F*_*ST*_. The rationale under this split is that because core genes are more differentiated between populations than random or peripheral genes, we should expect them to predict better those traits with a similar structuration behavior and *vice versa*. We found that traits above the 99^*th*^ percentile of the *F*_*ST*_ are systematically better predicted than less differentiated traits. However, we did not find significant differences between gene groups once the difference between traits was taken into account.

## Discussion

Characterizing the way genes contribute to phenotypic variation could prove highly valuable to better understand the genetic architecture of complex traits. With the advent of omics data, a huge amount of information is nowadays becoming available to fill the gap between variations at the DNA and phenotype levels. It is by the use of gene expression data that the present study aimed at gaining insights into the genetic architecture behind complex traits.

One key premise in the study was the availability of a common garden experiment comprising relevant samples of natural variation, in our case black poplar from Western Europe. Such an experimental setting makes it possible to accurately evaluate phenotypes to calibrate and serve as a target for predictions. Indeed, evaluating all the genotypes in a given location with experimental design and replicates enabled to unravel the confounding between genotype and macro-environment (or micro-environment) that typically occur when considering genotypes in the wild (de Villemereuil et al., 2016). Likewise, RNAseq data were collected on up to two biological replicates in the common garden and also corrected for environmental and design covariables, to obtain the genotypic BLUP, which is the genetic value of the genotype. Such adjustments at both phenotypic and genomic ends provided proper grounds with reasonable confidence in the absence of confounding effects for the study of associations between the two sources of data.

Two recent works used RNAseq in natural populations of plants to build co-expression networks and study the relationship between network topology and patterns of natural selection (Josephs et al., 2017; Mähler et al., 2017). While they found differences in natural selection among genes given their connectivity within networks, they did not investigate how these differences affect phenotypic variation. We thus embraced the classic WGCNA approach (Langfelder and Horvath, 2008) to build the co-expression network within our dataset in order to study the relationship between gene connectivity and phenotypic prediction. This clustering of genes gave us different groups that we found to be differently correlated to traits values and according to sites. However, this method was simply for us a way to obtain a centrality score for each gene, with the subsequent possibility to classify them into core and peripherals. The biological interpretation of correlations between gene groups and traits would clearly deserve further work which is beyond the scope of the present study. We based our definition of core and peripheral on Mähler et al. (2017), as respectively the 10% most central and most peripheral genes. The only specificity of our work here is that we did not discard, as it is classically done (called pruning in the WGCNA manual), the genes from the grey group, i.e. those showing a poor membership to any other module. We considered instead two alternative peripheral sets by keeping or excluding genes from the grey group. The pertinence of kME as a classification criterion became evident in our study when looking at the differences between core and peripherals in terms of classic quantitative and population genetic parameters. Core genes (high kME) showed high levels of population differentiation, mostly in quantitative genetic terms (*Q*_*ST*_), while being simultaneously less variable than the rest of the genes. Such results would suggest that core genes are genes potentially subjected to divergent selection, with subsequently reduced levels of extant variation, and involved in local adaptations. Contrarily, peripherals (low kME) showed larger levels of variation with respect to their expression level and little structure across populations, suggesting less selection pressure or weaker connection to selected traits, with mostly stabilizing selection patterns across populations. Therefore, despite the fact that a subdivision in core and peripherals is somehow an oversimplification, an extreme contrast of an otherwise continuous phenomenon, it helped to reveal the different natures of genes characterized by extreme values of kME.

To further test whether this gene categorization matters for trait prediction, we decided to go one step further by trying to predict traits from the different gene sets. We also wanted to have a gene set designed to be composed of good predictors of the traits. We thus used the Boruta algorithm (Kursa and Rudnicki, 2010) to select genes, by performing random forest predictions and selecting the genes with the highest prediction importances. We have to keep in mind that random forest algorithm allows for implicit interactions between predictors (here gene expressions, (McKinney et al., 2006; Chen et al., 2007; Jiang et al., 2009)). Results pinpointed again one feature differentiating the behavior of core and peripheral genes. Cores were largely overrepresented in the different Boruta selections (by at least 38% of Boruta genes), involving systematically the same 114 genes across all threshold p-values (153 over 3 values). Peripherals were systematically underrepresented to a very large extent (less than 7%). Although the remaining genes, neither cores nor peripherals according to our previous definition, were the majority (53%) among the selected by Boruta, they were sampled from a vaster pool of more than 27,000 genes. Another important result from the Boruta selections is the fact that relaxing the p-value threshold (from 0.01 to 0.2) did not increase the size of the resulting selection set, while the set itself could change partially in composition across different thresholds. One can assume that relaxing the threshold would lead to increasing the number of features if these acted independently and contributed with novel information. The fact that numbers did not change substantially, while the composition was indeed impacted, leads to thinking that features are deeply interconnected and do not add up independently. This would suggest that different arrangements of genes could contain comparable levels of information or, in other words, that genes bear some redundancy through networks of interactivity.

With these 6 genes sets, we predicted 17 phenotypic traits with 2 alternative algorithms, one expected to capture mostly additivity between predictors (LM), the other one interactivity (NN). As expected, the full set resulted in best predictions with the LM model (NN not available), as it comprised all available genetic information. Core genes, however, were far from being the best set to predict the different traits under either of the two algorithms. Such results would be a priori surprising considering previous statements on the composition of Boruta selection where cores had an important contribution. The key difference, however, is that cores were not the only contributors to the Boruta sets. It seems that cores are able to summarize key information for quality predictions but require a complementary contribution from other interacting genes to round up the optimal set. This is better reflected by the performance of the Boruta set, which obtained the best performance predicting traits under the NN algorithm. To some extent, the NN algorithm exploits the interactivity between features (genes) already present in the Boruta set, itself obtained through the random forest heuristics that are particularly sensitive to interactions. To some extent, the high connectivity of high kME value core genes is well captured by interaction sensitive algorithms to improve prediction.

In a contrasting way, Boruta and core sets performed poorly under LM modeling, where the two classes of peripherals obtained the best predictabilities. Such a performance from peripherals is somehow surprising, in the sense that this class of genes, notably the grey module, is usually pruned from transcriptomic studies, while they seem nonetheless to harbor important biological information that is relevant to the trait variation. Judging from the nature of the LM modeling, peripherals would have more a type of additive gene action, which could be in turn a penalizing feature when a reduction in the number of genes operates to focus only on the most relevant ones. Thus, peripherals appear to be relevant when allowed to contribute cumulatively to prediction, although they can be otherwise easily summarized by more integrative genes when variable selection procedures operate to obtain optimal sets. It is important to note, however, that adding peripherals (following an increasing kME) beyond the numbers present in their original sets did not improve predictability (**Figure S10**), suggesting the existence of a plateau in their capacity to explain trait variation. The low connectivity of peripheral genes, reflecting independent features, is best exploited by linear model approaches capturing mostly additive genetic actions.

Finally, random sets offered a convenient framework for inferences when comparing gene sets. Their performance in terms of predicting quality was never the best under either of the alternative modeling approaches (LM or NN) but was good enough to suggest that relevant information can be nevertheless obtained from many different gene sets, pointing at some degree of pervasive redundancy in the genetic architecture of traits. In practical terms, when a trait prediction is required but there is no biological *a priori* on the choice of genes, a random set modeled through LM appears like a satisfying solution. This is not far from the SNP based counterpart in genome-wide evaluation (Meuwissen et al., 2001), where markers are often a choice that is not driven by biological context. However, if some previous selection of genes is required, the combination of Boruta selection and subsequent NN modeling has been shown here to be a good option for predictability on a reduced genic panel. Indeed, Boruta is an advantageous alternative in genomic evaluation for breeding to more classic methods, often based on the imposition of a priori constraints for shrinkage or variable selection (de los Campos et al., 2013).

One of the particularities of core genes, that of showing highly structured genetic variation among populations, led us to think that they might be preferentially involved in traits also showing high levels of *Q*_*ST*_. Such a hypothesis was not confirmed by our results, where highly structured traits were generally better predicted than traits with no apparent structure, but with no clear differences in such an advantage between gene sets. Therefore, the highly structured core genes did not contribute to improving the prediction of highly structured traits, suggesting that trait covariation between populations is affected by other genic sources not conveniently unraveled here. It is important to note that prediction quality is highly variable between traits, somehow masking the differences that might be found between gene sets. We have already pinpointed the relevance of kME in establishing a gradient of genes whose extremes show different behaviors in quantitative and population genetics statistics. These extremes also contribute differently to the explanation of phenotypic variability, through the light of different prediction models. One aspect that remained unanswered, however, is to what extent kME is also relevant to prediction without circumscribing our scope to the extremes. When computing the correlations between connectivity (kME) and prediction coefficients (importance in terms of effect) from LM across all the full set of genes, results showed that there are some strong positive correlations for three of the traits (Circ.Orl, S.G.Orl and Extractives.Orl). However, there is not a systematic trend across all the traits, suggesting that other differences in their genetic variability and genomic architectures might be also of importance here.

In the end, differential connectivity as reflected by our kME gradient from gene expressions pinpoints at the importance of mechanisms of gene interactions in the genetic architecture of traits. On top of the DNA sequence, the superposing layer of transcriptomics adds up the intermediate pattern of gene interactions and physiological epistasis, before the final level of phenotypic expression (Schrag et al., 2018). It is important to note, however, that such gene interaction at the transcriptomic level is not directly or necessarily related to epistasis in the context of statistical genetics literature, i.e. the interaction effect between alleles from different loci on a given phenotype (Cordell, 2002). The extent to which connectivity or transcriptomic interactivity relates to that level of epistasis is beyond the scope of current work but clearly deserves further investigation.

## Conclusion

This work shows that all genes seem important to some extent to predict phenotypes. If the Boruta selection leads us to think that core genes may be very important, prediction results across a range of phenotypes underlined that they are not the only ones. The information that they contain has to be completed by other genes. The mean connectivity score (kME) of the Boruta sets is around 0.7. However, as genes seem to be very interactive, predicting a phenotype with a subset of genes summarizing the information is possible and efficient. Our work is globally in accordance with the recent work on the omnigenic model (Boyle et al., 2017; Liu et al., 2019), describing that all genes expressed in an organ participate in the traits of that organ. We are also able to predict phenotypes of an organ or at the organism level, with gene expression from another organ. However predicting and explaining are 2 different things, and the information contained by genes may be too redundant to lead us to good mechanistic models from statistical ones. Statistical models may, nevertheless, provide information on the ranking of importance of the genes involved in a phenotype.

## Materials and Methods

### Samples collection

As described in previous works (Gebreselassie et al., 2017; Guet et al., 2015), we established in 2008 a partially replicated experiment with 1160 cloned genotypes, in two contrasting sites in central France (Orléans, ORL) and northern Italy (Savigliano, SAV). At ORL, the total number of genotypes was 1,098 while at SAV there were 815 genotypes. In both sites, the genotypes were replicated 6 times in a randomized complete block design. At SAV, the trees were pruned at the base after one year of growth (winter 2008-2009) to remove a potential cutting effect and were subsequently evaluated for their growth and wood properties during winter 2010-2011. At ORL, the trees had the same pruning treatment after two years of growth (winter 2009-2010) and were also subsequently evaluated for growth and wood properties after two years (winter 2011-2012). After evaluation, we pruned again for a new growth cycle. In their fourth year of growth of this third cycle (2015), 241 genotypes present in two blocks of the French site were selected to perform sampling for RNA sequencing. In the end, we obtained transcriptomic data from 459 samples, 218 genotypes duplicated in the two blocks and 23 genotypes available from only one block. These 241 genotypes were representative of the natural west European range of P. nigra through 11 river catchments in 4 countries (**Table S3**).

We described 14 of the 17 phenotypic traits in previous work (Gebreselassie et al., 2017). Briefly, these traits can be divided into two categories, growth traits and biochemical traits which were all evaluated on up to 6 clonal replicates by genotype at each site after two years of growth in the second cycle. The first set is composed of the circumference of the tree at a 1-meter height measured in Savigliano at the end of 2009 (CIRC2009.Sav) and in Orléans at the end of 2011 (CIRC2011.Orl). The second set is composed, each time at both sites, of measures of ratios between the different components of the lignin, p-hydroxyphenyl (H), guaiacyl (G) and syringyl (S) (H.G.Orl, H.G.Sav, S.G.Orl and S.G.Sav), measures of the total lignin content (Lignin.Orl : measure of the lignin in Orléans, Lignin.Sav: measure of the lignin in Savigliano), measure of the total glucose (Glucose.Orl and Glucose.Sav), measure of ratio between 5 and 6 carbon sugars (C5.C6.Orl and C5.C6.Sav) and measure of the extractives (Extractives.Orl and Extractives.Sav). For each of these traits, we computed mean values per genotype previously adjusted for microenvironmental effects (block or spatial position in the field).

The 3 remaining traits were measured in 2015 on the trees harvested for the RNA sequencing experiment (2 replicates per genotype). They include the mean diameter of the stem section harvested for RNA sequencing (MeanDiameter), the date of bud flush of the tree in 2015 (Date3Doy) and the basic density of the wood (Infraden). Date of bud flush consisted of a prediction of the day of the year at which the apical bud of the tree was in stage 3 according to the scale defined in Dillen et al. (2009). Predictions were done with a lowess regression from discrete scores recorded at consecutive dates in the spring of 2015. Wood’s basic density was measured on a piece of wood from the stem section harvested for RNA sequencing following the Technical Association of Pulp and Paper Industry (TAPPI) standard test method T 258 “Basic density and moisture content of pulpwood”.

### Transcriptome data generation

We sampled stem sections of approximately 80 cm long starting at 20 cm above the ground and up to 1 meter in June 2015. The bark was detached from the trunk in order to scratch young differentiating xylem and cambium tissues using a scalpel. The tissues were immediately immersed in liquid nitrogen and crudely ground before storage at −80°C pending milling and RNA extraction. Prior to RNA extraction, the samples were finely milled with a swing mill (Retsch, Germany) and tungsten beads under cryogenic conditions with liquid nitrogen during 25 seconds (frequency 25 cps/sec). About 100 mg of milled tissue was used to isolate separately total RNA from xylem and cambium of each tree with RNeasy Plant kit (Qiagen, France), according to manufacturer’s recommendations. Treatment with DNase I (Qiagen, France) to ensure the elimination of genomic DNA was made during this purification step. RNA was eluted in RNAse-DNAse free water and quantified with a Nanodrop spectrophotometer. RNA from xylem and cambium of the same tree were pooled in an equimolar extract (250 ng/*μ*L) before sending it to the sequencing platform.

RNAseq experiment was carried out at the platform POPS (transcriptOmic Platform of Institute of Plant Sciences - Paris-Saclay) thanks to IG-CNS Illumina Hiseq2000. RNAseq libraries were constructed using TruSeq Stranded mRNA SamplePrep Guide 150310 47 D protocol (Illumina®, California, U.S.A.). The RNAseq samples have been sequenced in single-end reads (SR) with an insert library size of 260 bp and a read length of 100 bases. Images from the instruments were processed using the manufacturer’s pipeline software to generate FASTQ sequence files. Ten samples by lane of Hiseq2000 using individually barcoded adapters gave approximately 20 millions of SR per sample. We mapped the reads on the *Populus trichocarpa* v3.0 transcriptome with bowtie2 (Langmead and Salzberg, 2012), and obtained the read counts for each of the 41,335 transcripts by homemade scripts (a median of 17 millions of reads were mapped per sample, with a minimum of 6 and a maximum of 42 million). Initially, we considered using the genotype means to reduce our data volume. However, differences between replicates were not normally distributed, because of variation in gene expression due to plasticity. We thus could not summarize our data with their mean, as it would have removed this information and finally we chose to keep replicates as separate data samples.

### Filtering the non-expressed genes, normalization and variance stabilization

We started cleaning our raw count data by removing the transcripts without at least 1 count in 10% of the individuals. From the original 41,335 genes, 7,106 were thus removed, leaving 34,229 genes. After this first filtration, we normalized the raw count data by Trimmed Mean of M-values (TMM, edgeR (Robinson and Oshlack, 2010)). As most features are not differentially expressed, this method takes into account the fact that the total number of reads can be strongly influenced by a low number of features. Then, we calculated the counts per millions (CPM (Law et al., 2014)).

To stabilize the variance of the CPM data, we computed a *log*_2_(*n* + 1) instead of a *log*_2_(*n* + 0.5) typically used in a voom analysis (Law et al., 2014), to avoid negative values, which are problematic for the rest of the analysis.

### Computing the BLUP, heritability, and *Q*_*ST*_ while correcting the co-variables

As the sampling ran along 2 weeks, we expected environmental variables to blur the signal. To understand how our data were impacted, we ran a PCA analysis to identify the impact of each cofactor (**Figure S1**). We identified the block and the sampling date and time as cofactors with a substantial impact.

A 12k bead chip (Faivre-Rampant et al., 2016) provided 7,896 SNPs in our population. A genomic relationship matrix between genotypes was computed with these SNPs with LDAK (Speed et al., 2012), and further split into between (mean population kinship, **K**_**b**_) and within-population relationship matrices (kinship kept only for the members of the same population, all the others are equal to 0, **K**_**w**_). These matrices were used in a mixed linear model to compute the additive genetic variances between and within populations for the expression of each gene:

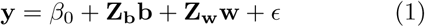

Where, **y** is a gene expression vector across individual trees, *β*_0_ is a vector of fixed effects (overall mean or intercept); **b** and **w** are respectively random effects of populations and individuals within populations, which follow normal distributions, centered around 0, of variance 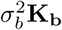 and 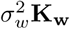. *σ*_*b*_ and *σ*_*w*_ are the between and within-population variance components and **K**_**b**_ and **K**_**w**_ are the between and within-population kinship matrices. **Z**_**b**_ and **Z**_**w**_ are known incidence matrices between and within populations, relating observations to random effects **b** and **w**. *E* is the residual component of gene expression, following a normal distribution centered around 0, of variance 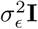, where *σ*_*ϵ*_ is the residual variance and **I** is an identity matrix.

We used the function “remlf90” from the R package breedR (Muñoz and Sanchez, 2017) to fit the model, with the Expectation-Maximization method followed by one round with Average-Information algorithm to compute the standard deviations. From the resulting between and within-population variance components, we computed the best linear unbiased predictors of between and within population random genetic effects (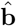 and **ŵ**, respectively) and summed them up to obtain the total genetic value for each gene expression (*BLUP*). We also computed heritability (*h*^2^) and population differentiation estimates (*Q*_*ST*_) for each gene expression as follows:

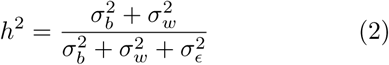

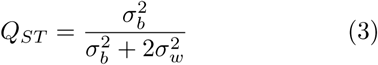

Finally, we computed for each gene expression the coefficient of genetic variation (*CV*_*g*_) by dividing its total genetic variance 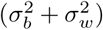 by its expression mean.

### Other population statistics

We further used a previously developed bioinformatics pipeline to call SNPs within our RNA sequences (Rogier et al., 2018). Briefly, this pipeline involves classical cleaning and quality control steps, mapping on the *P. trichocarpa* v3.0 reference genome, and SNP calling using the combination of four different callers. We ended up with a set of 874,923 SNPs having less than 50% of missing values per genotype. The missing values were further imputed with the software FImpute (Sargolzaei et al., 2014). We validated our genotyping by RNA sequencing approach by comparing the genotype calls with genotyping previously obtained with an SNP chip on the same individuals (Faivre-Rampant et al., 2016). Genotyping accuracy based on 3,841 common positions was very high, with a mean value of 0.96 and a median value of 0.99. The imputed set of SNP was then annotated using Annovar (Wang et al., 2010) in order to group the SNPs per gene model of *P. trichocarpa* reference genome. For each SNP, we computed the overall genetic diversity statistics with the hierfstat R package (Goudet and Jombart, 2015) and this statistic was then averaged by gene model in order to get information on the extent of diversity. We further computed *PCadaptscore* with the pcadapt R package (Luu et al., 2017) with 8 retained principal components. Here again, *PCadapt scores* were then summarized (averaged) by gene-model in order to get information about their potential involvement in adaptation. Based on the principal component analysis, pcadapt is more powerful to perform genome scans for selection in next-generation sequencing data than approaches based on *F*_*ST*_ outliers detection (Luu et al., 2017). We found a positive correlation between *F*_*ST*_ and *PCadapt score* (data not shown), but *PCadapt score* highlighted differences between Core, random and peripheral gene sets (Figure 3) while *F*_*ST*_ did not.

### Hierarchical and k-means clustering

We performed a weighted correlation network analysis with the R package WGCNA (Langfelder and Horvath, 2008) on our full RNAseq gene set. We followed the classic approach, except that we first ranked our expression data, to work subsequently with Spearman’s non-parametric correlations and avoid problems due to linear modeling assumptions. We first chose the soft threshold with a power of 12, which is the classical value for signed networks (and default value in WGCNA) (*R*^2^ = 0.81, connectivity: mean = 195.17, median = 9.23, max = 1403.96, **Figure 2A**). Then, we used the automatic module detection (function “blockwiseModules”) via dynamic tree cutting with a merging threshold of 0.25, a minimum module size of 30 and bidweight midcorrelations (**Figure 2B**). All other options were left to default. This also computes module eigengenes.

To sort the traits, we clustered their scaled values with the pvclust R packages (Suzuki and Shimodaira, 2015), the Ward agglomerative method (“Ward.D2”) on correlations (**Figure 2B, 2C, Figure S2**). The clustering on euclidean distance results in the exact same hierarchical tree. Correlations between traits and gene expression or module eigengenes were computed as Spearman’s rank correlations (**Figure 2B, 2C**). We also performed a k-means clustering with the R package coseq (Rau and Maugis-Rabusseau, 2017) considering 10 initial runs, 1000 iterations, without any other data transformation, and for a number of clusters (K) between 2 and 20. At first, it identified a K without strong agreement between the two evaluation algorithms included in coseq. We thus further computed additional rounds of k-means clustering, around the previously identified K (plus or minus 5 clusters), with 100 initial runs and 10000 iterations, until both evaluation algorithms agreed.

### Machine learning

#### Boruta gene expression selection

In addition to the inconvenience of working with a large number of features (time and power consumption), most machine learning algorithms perform better when the number of predicting variables used is kept as low as the optimal set (Kohavi and John, 1997). We thus performed an all relevant variable selection (Nilsson et al., 2007) with the Boruta function (Kursa and Rudnicki, 2010) from the eponym R package, with 4 p-value thresholds (1, 5, 10 and 20%), on the training subpart of the full gene expression set, for each phenotype independently. Then, features that were not rejected by the algorithm were pooled together, so that all the important genes were in the selected gene pool, one pool for each p-value threshold. The enrichment in core or peripheral genes in each of these pools was evaluated by Fisher’s exact test for count data (“fisher.test” function in the stats R package).

#### Models

Both additive linear model (ridge regression) and interactive neural network models were computed by the R package h2o (LeDell et al., 2019). They both used the gene expression sets as predictors and one phenotypic trait at a time as a response. Gene sets were split by the function “h2o.splitFrame” into 3 sets, a training set, a validation set and a test set, with the respective proportions of 60%, 20%, 20%. We checked that the split preserves the distribution of samples within populations. The training set was used to train the models, the validation set was used to validate and improve the models, while the test set was used to compute and report prediction accuracies as *R*^2^ between observed and predicted values within this set and using the function “R2” of the R package compositions (van den Boogaart et al., 2018). This set has never been used to improve the model and therefore represents a proxy of new data, avoiding the report of results from overfitted models. All the reported predictions scores were computed on this test set. These results are thus representing reallife predictions and are not subject to over-fitting.

For linear models, we used the function “h2o.glm with” default parameters, except 2-folds cross-validation and alpha set at zero to perform a ridge regression. The same splits and score reporting methods were used.

Neural networks have the reputation to be able to predict any problem, based on the Universal approximation theorem (Cybenko, 1989; Hornik et al., 1989). However, this capacity comes at the cost of a very large number of neurons in one layer, or a reasonable number of neurons per layer in a high number of layers. Both settings lead to difficult interpretation when very many gene expressions are involved. In that sense, we chose to keep our models simple, with two layers of a reasonable number of neurons. This obviously comes at the price of lower prediction power. However, we believe that these topologies give us the power to model 2 levels of interactions between genes (1 level per layer). Furthermore, since both methods yielded comparable prediction *R*^2^ (median ridge regression *R*^2^ = 0.19, mean neural network *R*^2^ = 0.173), this complexity seemed appropriate. To find the best models for neural networks, we computed a random grid for each response. We tested the following four hyperparameters: (i) activation function (“Rectifier”, “Tanh”, “Rectifier-WithDropout” or “TanhWithDropout”); (ii) network structure; (iii) input layer dropout ratio (0 or 0.2) (iv) L1 and L2 regularization (comprised between 0 and 1 ×10^−4^, with steps of 5 ×10^−6^). Network structure corresponded to the number of neurons within each of the two hidden layers, which was based on the number of input genes (*h*). The first layer was composed of *h*, 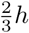 or 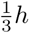 neurons. The second layer had a number of nodes equal or lower to the first one and is also composed of *h*, 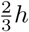 or 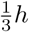 neurons. This represented a total of 6 different structures. We performed a random discrete strategy to find the best search criteria, computing a maximum of 100 models, with a stopping tolerance of 10^−3^ and 10 stopping rounds. Finally, “h2o.grid” parameters were the following: the algorithm was “deeplearning”, with 10 epochs, 2 fold cross-validation, maximum duty cycle fraction for scoring is 0.025 constraint for a squared sum of incoming weights per unit is 10. All other parameters were set to default values. The best model was selected from the lowest RMSE score within the validation set.

## Declarations

## Availability of data and materials

This RNAseq project has been submitted to the international repository Gene Expression Omnibus (GEO) from NCBI (accession number: GSE128482). All steps of the experiment, from growth conditions to bioinformatic analyses are detailed in CATdb (Gagnot et al., 2007) according to the MINSEQE “minimum information about a high-throughput sequencing experiment”. Raw sequences (FASTQ) are being deposited in the Sequence Read Archive (SRA) from NCBI. Information on the studied genotypes is available in the GnpIS Information System (Steinbach et al., 2013).

## Authors’ Contributions

AC, LS, and VS designed the experiment, discussed the results and wrote this manuscript. AC ran the in silico experiment. MCL, VB, CPL, LT, MLMM, and VS contributed the RNAseq data production and analysis. VJ, OR and VS contributed to the SNP data production and analysis. MLMM and JCL contributed to the discussion on the methodology employed. All the authors read and approved this manuscript.

## Acknowledgements

The authors gratefully acknowledge the staff of the INRA GBFOR experimental unit for the establishment and management of the poplar experimental design in Orléans, the collection of wood samples in each site, and their contribution to phenotypic measurements on poplars in Orléans; Alasia Franco Vivai staff for management of the poplar experimental plantation in Savigliano, and M. Sabatti and F. Fabbrini for their contribution to phenotypic measurements on poplars in Savigliano. We acknowledge the staff of BioForA for their contribution to RNA collection in the field. We are grateful to the genotoul bioinformatics platform Toulouse Midi-Pyrennées for providing computing and storage resources. We would also like to thank M. Nordborg for useful discussions on this work and J. Salse for useful comments on the manuscript.

## Supplemental Material

### Supplemental tables

**Table S1:**
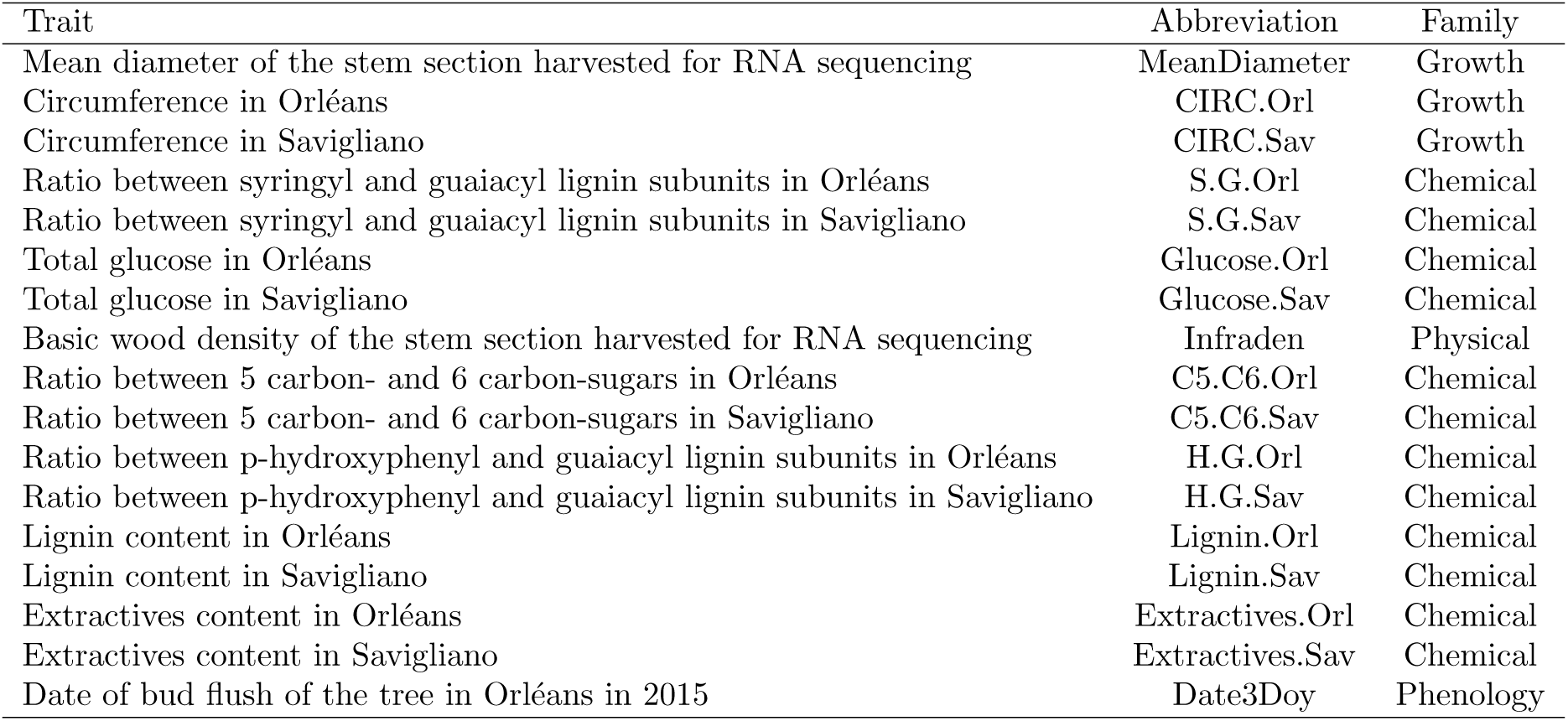
Correspondence between traits, their abbreviations, and families.

**Table S2:** Module membership of each gene (see **Supplemental file**).

**Table S3:** Number of genotypes sampled for each population.

| Population name | Country | Number of genotypes |
| --- | --- | --- |
| Adour | France | 36 |
| Basento | Italy | 5 |
| Dranse | France | 16 |
| Kuhkopf | Germany | 19 |
| Loire | France | 34 |
| NL | Netherlands | 4 |
| Paglia | Italy | 13 |
| Ramieres | France | 26 |
| Rhin | France | 15 |
| Ticino | Italy | 54 |
| ValAllier | France | 19 |

**Table S4:** Distribution of core and peripheral genes across modules.

| Module | core | peripheral | peripheral no grey |
| --- | --- | --- | --- |
| grey | 0 | 2942 | 0 |
| blue | 519 | 300 | 1438 |
| turquoise | 498 | 85 | 933 |
| yellow | 368 | 1 | 99 |
| brown | 258 | 13 | 152 |
| magenta | 119 | 12 | 167 |
| pink | 135 | 14 | 107 |
| red | 120 | 8 | 111 |
| lightcyan | 170 | 1 | 21 |
| cyan | 74 | 4 | 68 |
| salmon | 75 | 8 | 63 |
| grey60 | 136 | 1 | 1 |
| darkturquoise | 69 | 3 | 32 |
| purple | 45 | 4 | 52 |
| black | 70 | 5 | 21 |
| greenyellow | 56 | 3 | 20 |
| darkgrey | 37 | 1 | 21 |
| saddlebrown | 58 | 0 | 1 |
| violet | 53 | 0 | 0 |
| white | 27 | 0 | 23 |
| darkmagenta | 39 | 3 | 7 |
| lightyellow | 33 | 4 | 10 |
| orange | 45 | 0 | 0 |
| darkorange | 43 | 0 | 1 |
| darkred | 43 | 0 | 0 |
| royalblue | 37 | 0 | 5 |
| green | 25 | 0 | 14 |
| lightgreen | 37 | 0 | 0 |
| paleturquoise | 24 | 1 | 12 |
| skyblue | 32 | 1 | 4 |
| tan | 27 | 1 | 9 |
| darkgreen | 22 | 1 | 10 |
| darkolivegreen | 25 | 1 | 3 |
| midnightblue | 18 | 1 | 10 |
| steelblue | 25 | 1 | 3 |
| yellowgreen | 22 | 1 | 2 |
| sienna3 | 19 | 2 | 2 |
| skyblue3 | 19 | 0 | 0 |

### Supplemental figures

**Figure S1:**
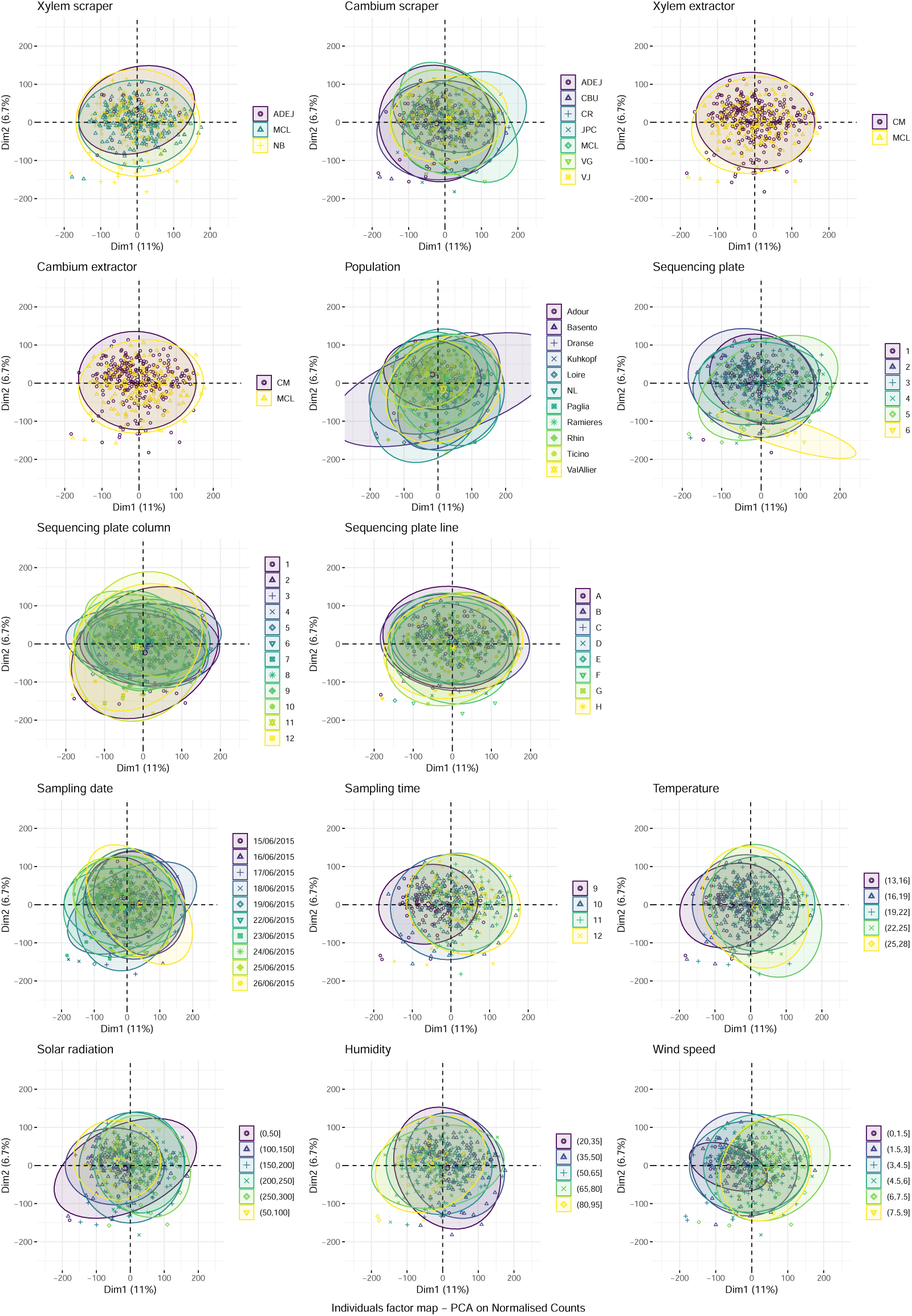
PCA of the different cofactors (Xylem and cambium scraper, extractor and extraction method, population, sequencing column, line and plate, the growth rate at harvest, sampling date, time, temperature, solar radiation, humidity and wind speed). Each of these represents the distribution of the individuals on the 2 first axes of the PCA (representing 17,7% of the variation), colored by class. Cofactors related to weather are presented in the 6 lower plots.

**Figure S2:**
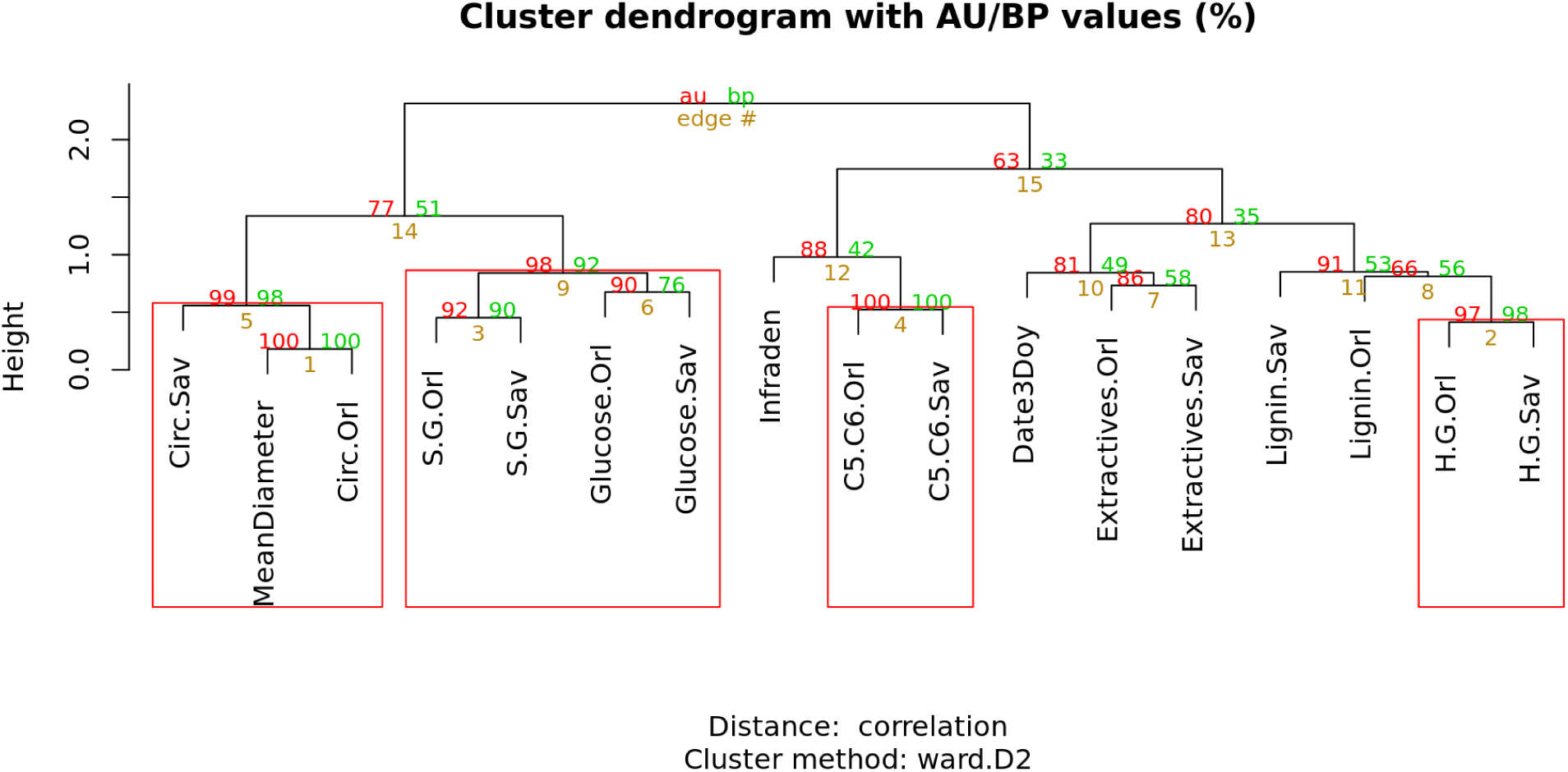
Scaled traits hierarchical clustering dendrogram computed from their correlations with Ward method (“Ward.D2”) by the R package pvclust. Approximately Unbiased (au, in red) and Bootstrap Probability (bp, in green) p-values indicated the degree of belief associated with clusters. Highly supported modules are framed by a red square, grouping (a) the mean sample diameter with the two circumference traits, (b) the S/G ratios with glucose composition, (c) the two C5/C6 together, and (d) the H/G ratios.

**Figure S3:**
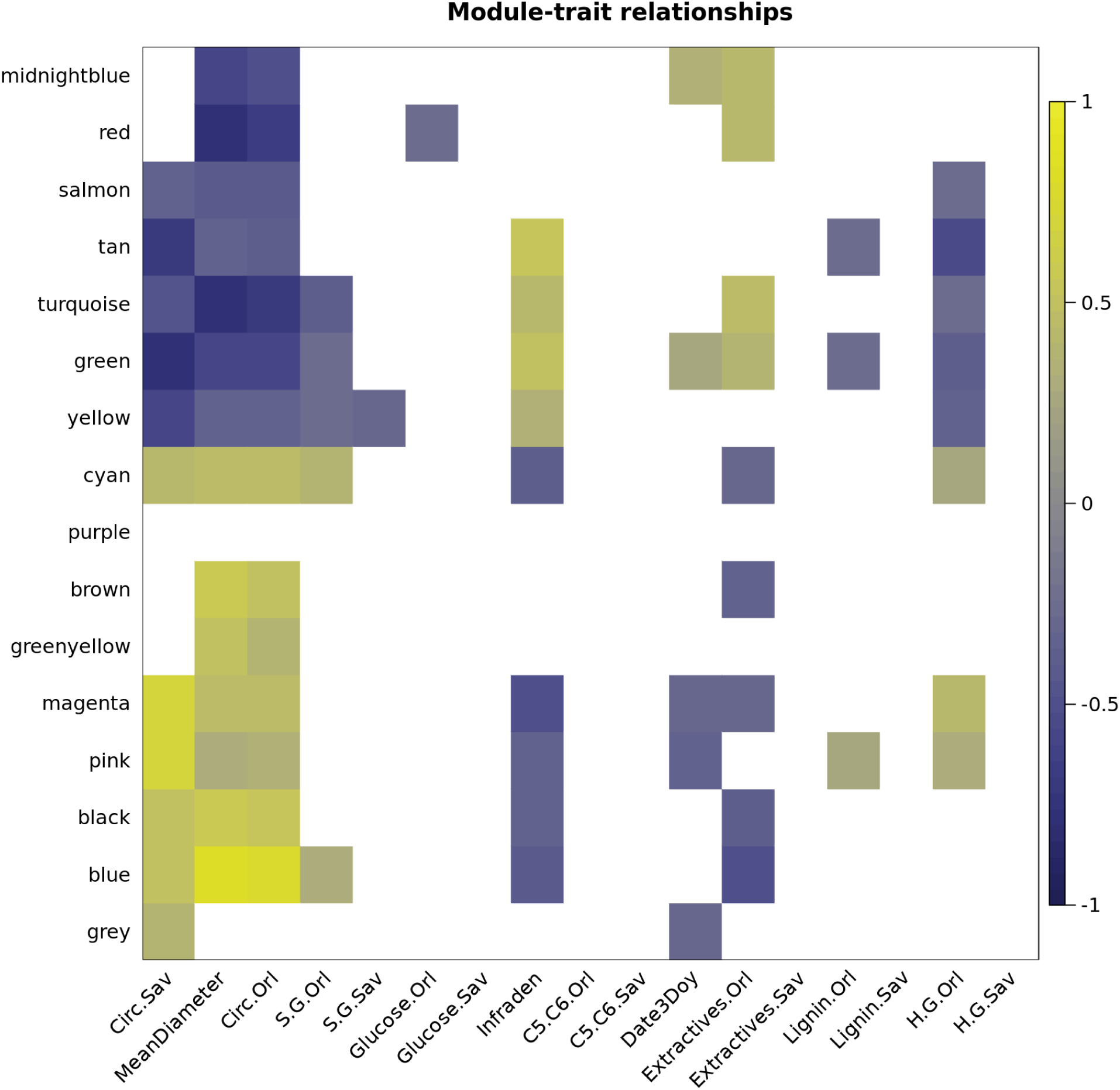
Heatmap of module-trait Spearman’s correlations, on a dark blue (high negative correlation) to light yellow (high positive correlation) scale. We removed correlations with a p-value lower than 5% after Bonferroni correction. From the total of 425 correlations, 72 remained.

**Figure S4:**
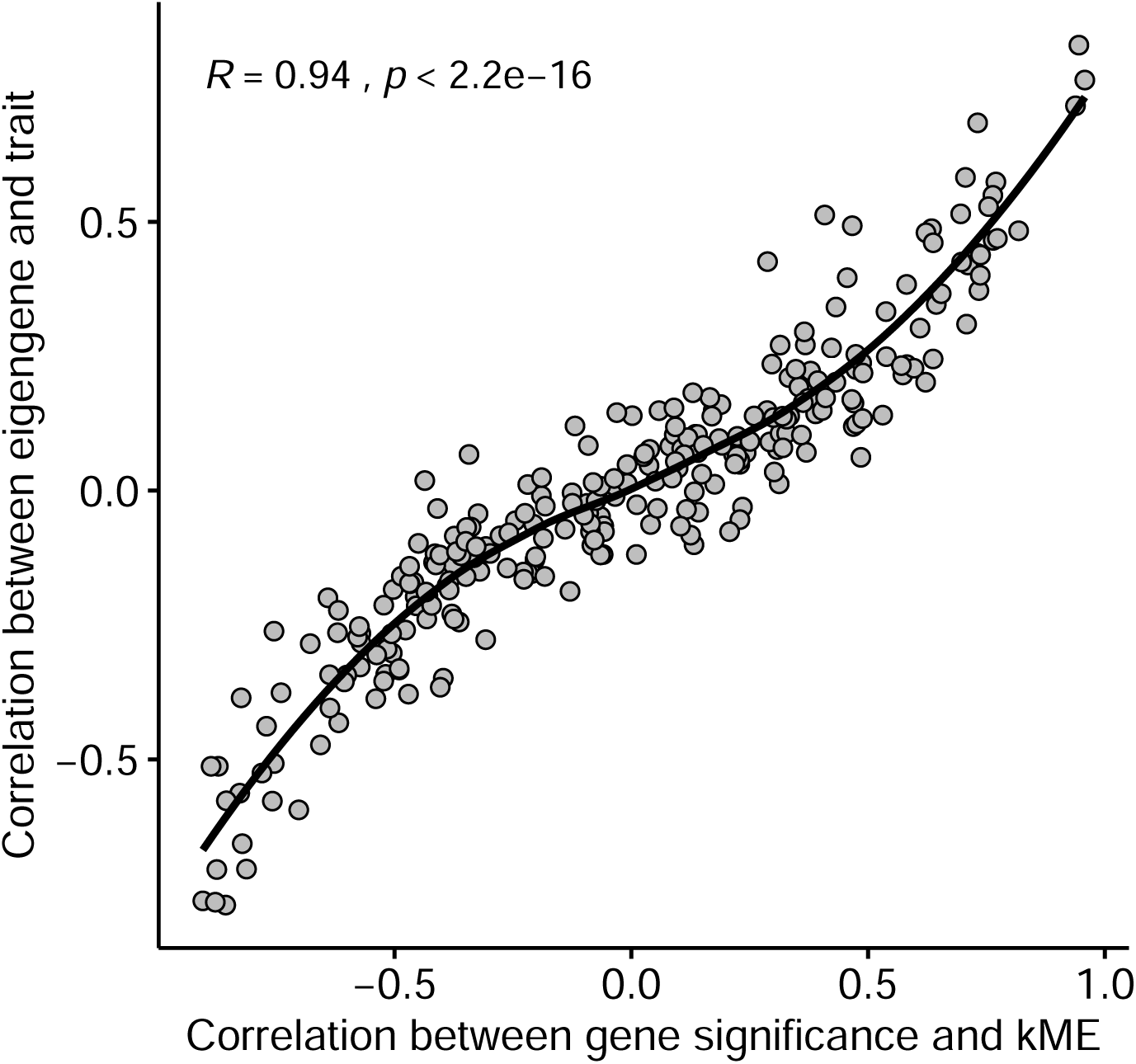
Relationship between Spearman’s correlations between module-trait (y-axis) and gene significance-kME (x-axis).

**Figure S5:**
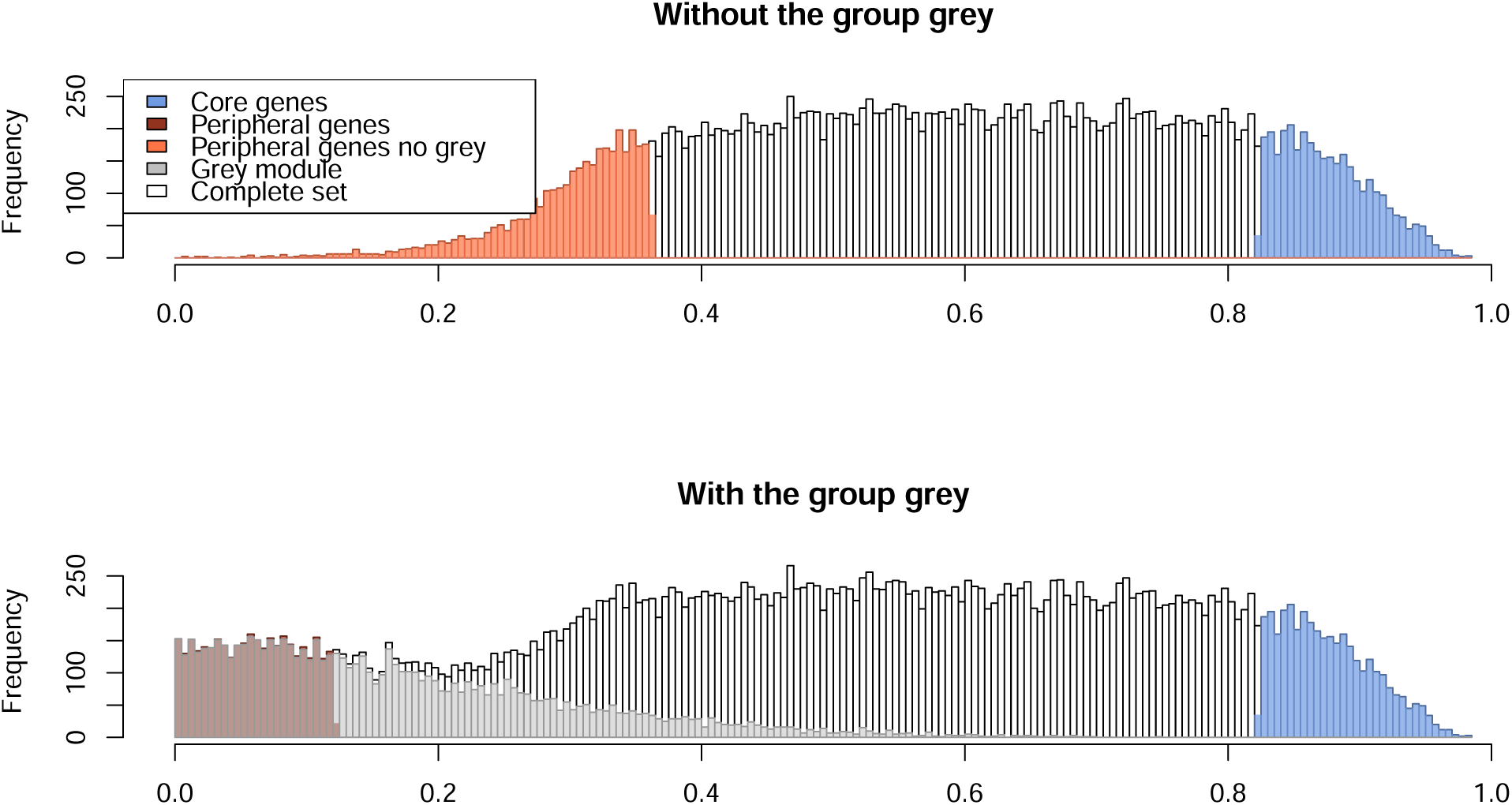
Histogram of the centrality scores without (top panel) or with (botom panel) the grey group. Core, peripheral and peripheral without grey sets are represented respectively by the blue, dark orange and orange bars. Random sets are distributed across the histogram and do not appear on this figure. Distribution of genes clustered in the grey module is represented by the grey bars, white bars are for other genes.

**Figure S6:**
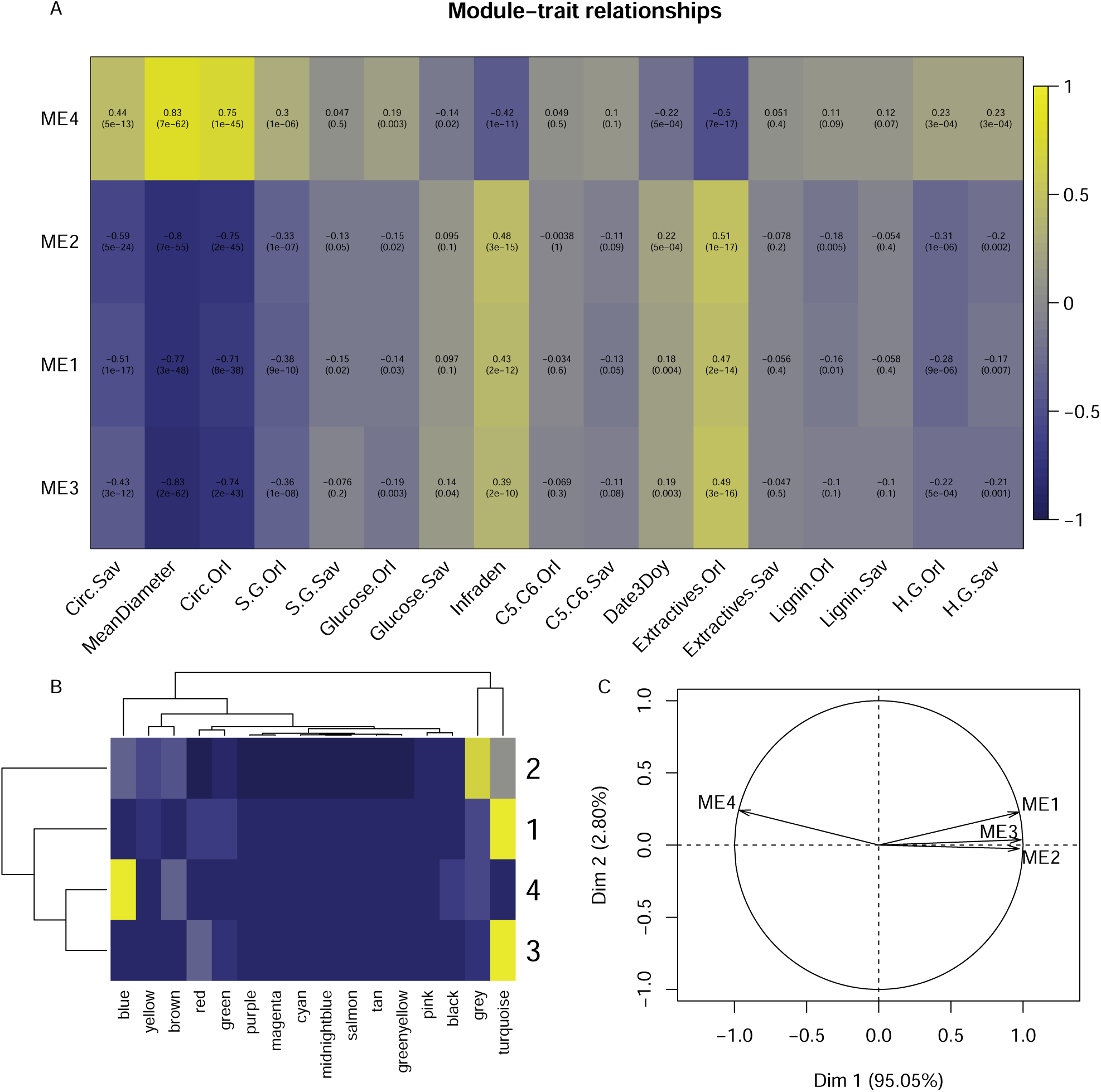
Gene expression k-means clustering (A) Correlation between eigengenes of modules identified by k-means clustering, on a light yellow (positive) to dark blue (negative) scale. P-values are indicated on the second line of each square. (B) Heatmap representing the concordance between WGCNA (abscissa) and k-means (ordinate) clusterings. (C) Principal component analysis graph of the k-means clustering.

**Figure S7:**
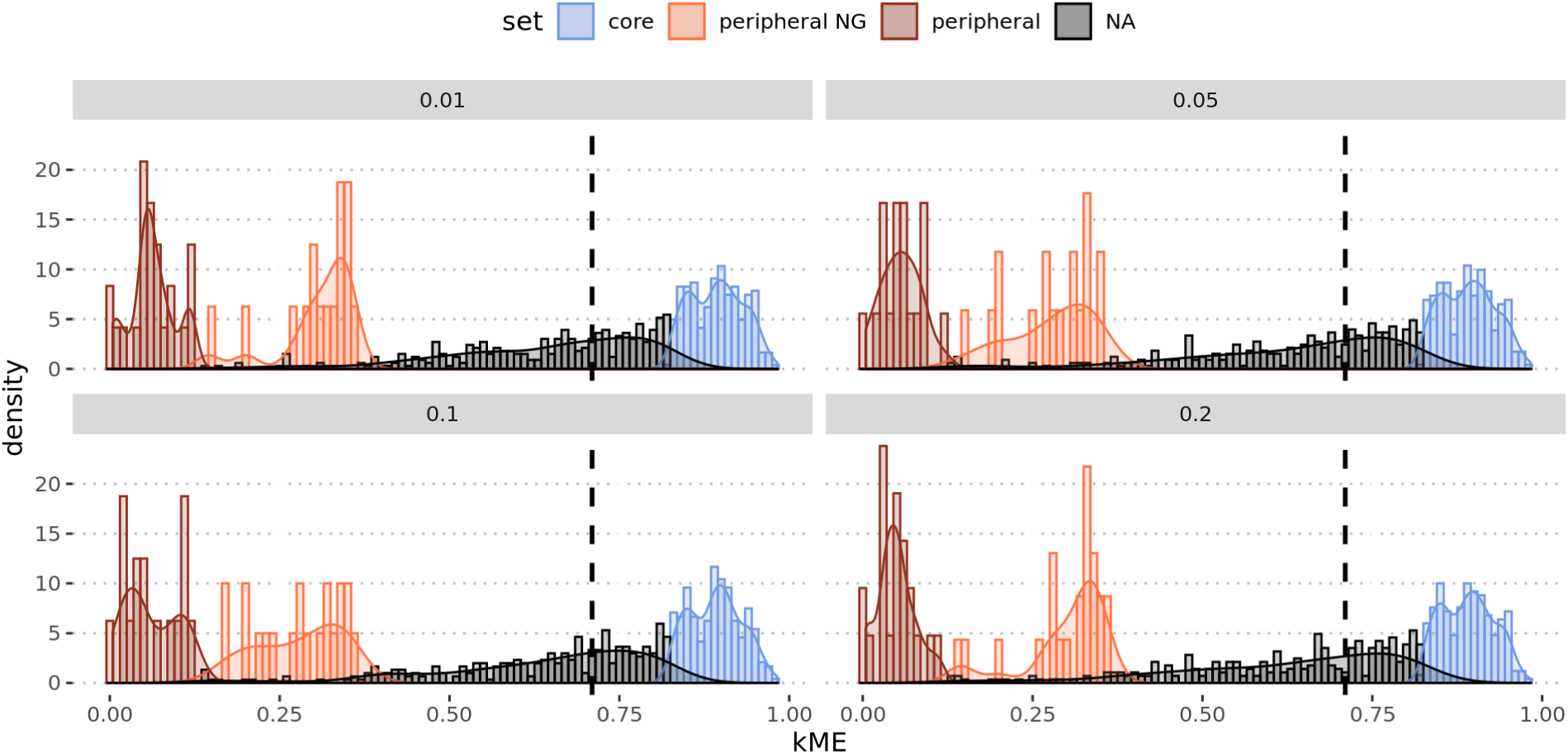
Distribution of the kME for the core (blue), peripheral NG (orange), peripheral (brown) and other (NA, in black) genes in the sets selected by Boruta for the different p-values (0.01, 0.05, 0.1 and 0.2).

**Figure S8:**
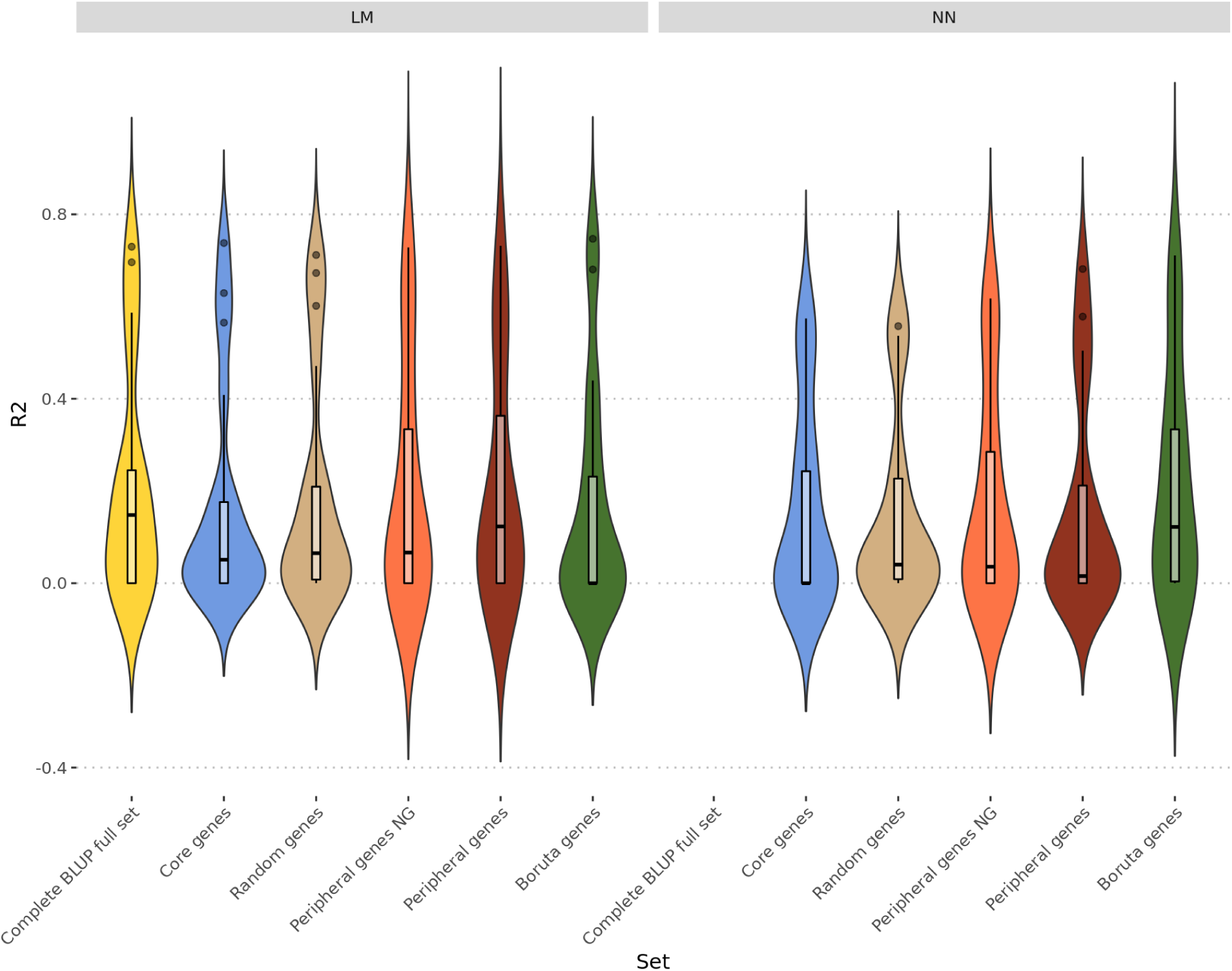
Violin and boxplots of prediction *R*^2^ across all phenotypes, split by model and gene sets.

**Figure S9:**
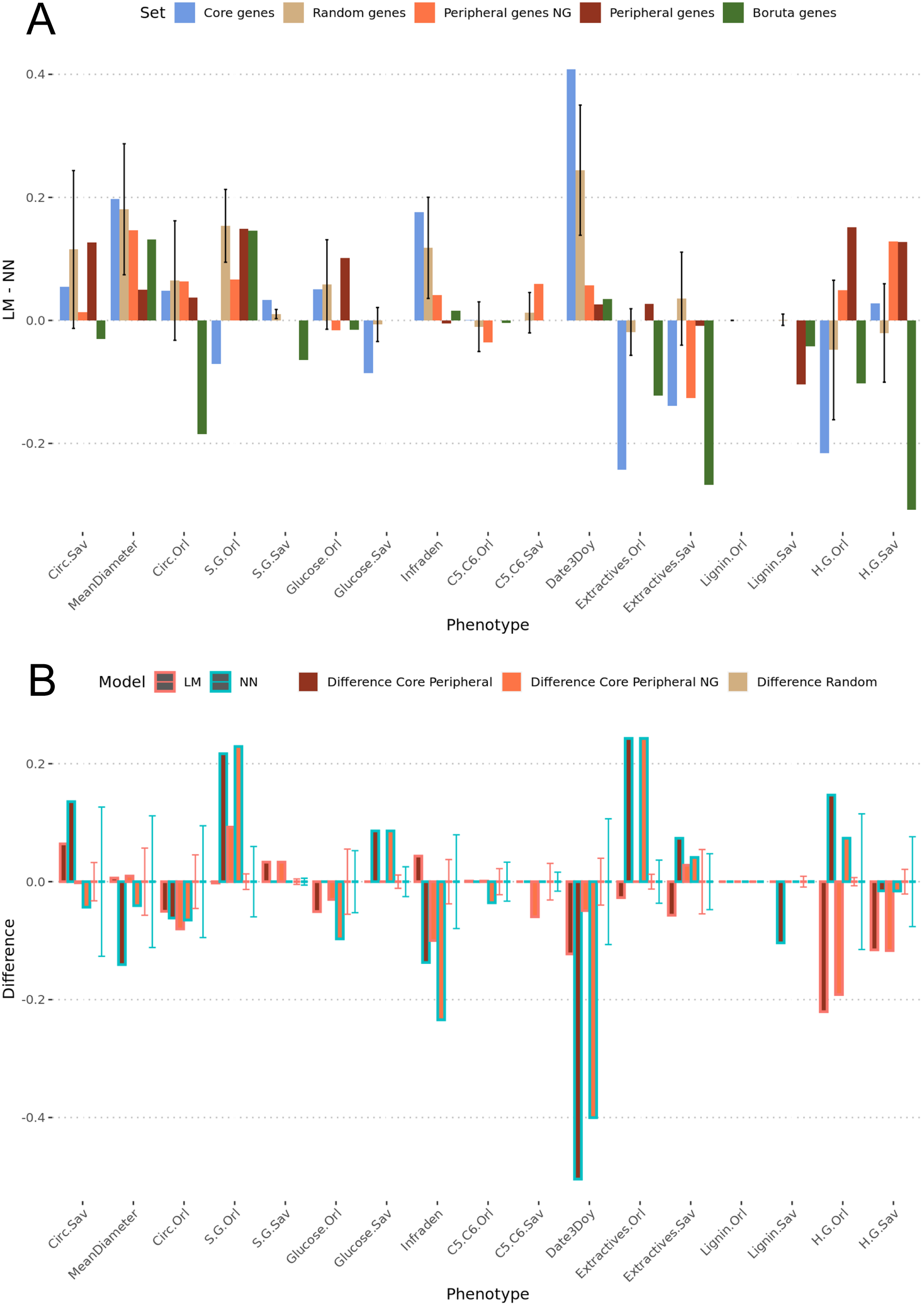
Difference of prediction scores (on the y-axis) between algorithms (A) and sets (B). (A) the difference between LM and NN prediction scores for the core (in blue), random (in grey), peripheral (in brown), peripheral (in orange) and Boruta gene sets (in green).(B) the LM differences are in red and the NN differences in turquoise and the color filling the bar represents the difference between core and peripheral genes in brown, core and peripheral NG in orange and between the random sets in grey. For the random pairs, error bars represent the first and third quartiles of the differences between pairs of randomized sets and the bar corresponds to the median.

**Figure S10:**
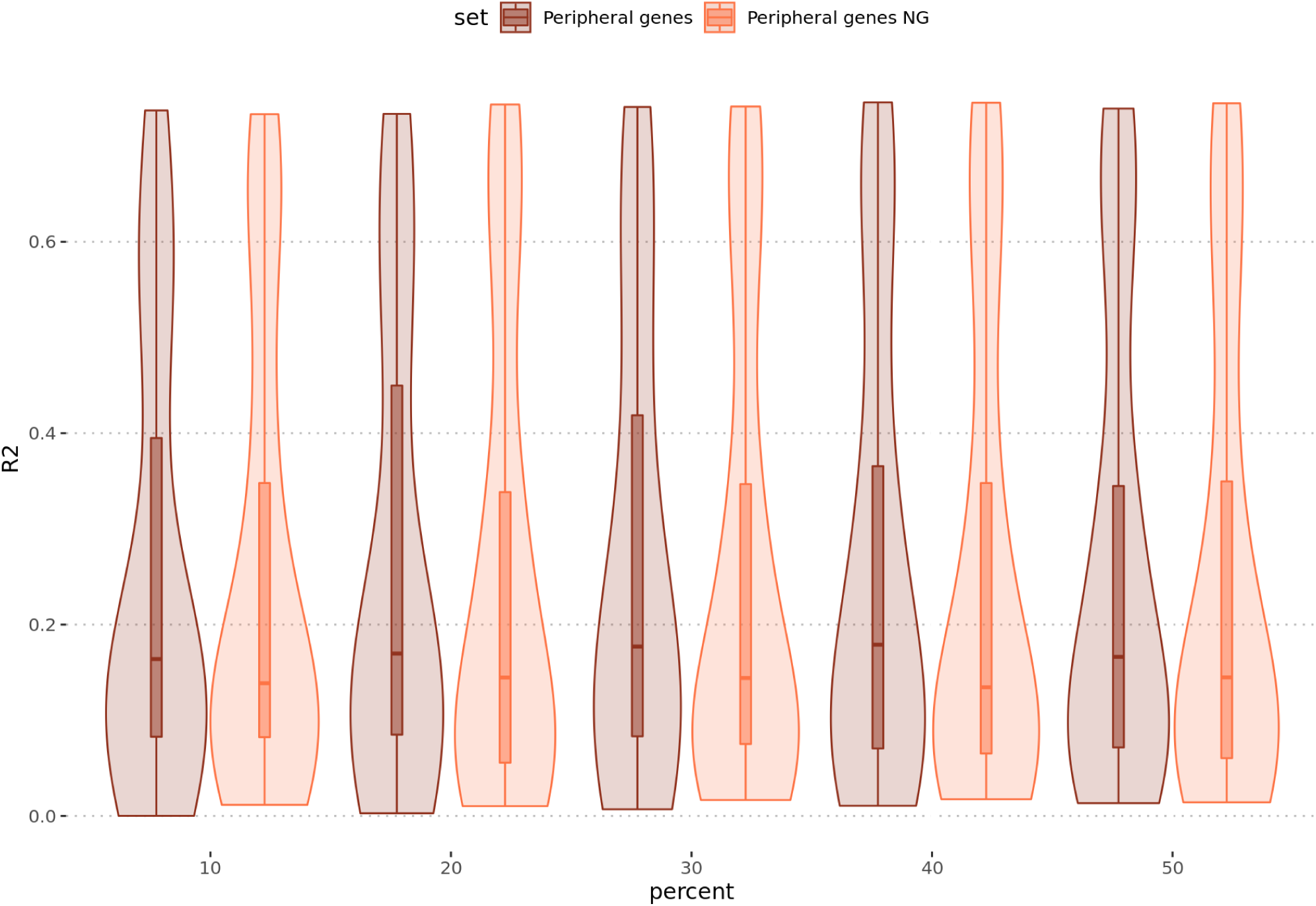
Violin plots of the predictions scores on test sets (*R*^2^ on the y-axis) for the LM Ridge algorithm for increasing sizes of the peripheral genes set (in brown) and the peripheral NG genes set (in orange), used for the predictions (in percent of the full set).

